# Lef1 Promotes Foxi1-Dependent Ionocyte Specification by Airway Submucosal Gland Progenitors

**DOI:** 10.64898/2026.10.13.758987

**Authors:** Weam Shahin, Javahir Hajibabazade, Olivia Snyder, Grace Fritz, Feng Yuan, Vitaly Ievlev, Thomas Lynch, Charles Yeaman, John F. Engelhardt

**Affiliations:** Department of Anatomy and Cell Biology, Roy J. and Lucille A. Carver College of Medicine, University of Iowa, Iowa City, IA 52242, USA

**Keywords:** Pulmonary ionocytes, airway submucosal gland myoepithelial cells (GMECs), ionocyte specification, Lef1, Foxi1, Bsnd, Cftr, anion conductance, 3D organoid swelling, airway regeneration

## Abstract

Pulmonary ionocytes are rare cells in submucosal gland (SMG)–containing airways thought to have specialized ion transport properties that regulate airway surface fluid and mucus composition. The origins of pulmonary ionocytes remain unclear, but their association with airway SMGs suggests they may arise from specialized progenitors in this compartment. We show that mouse glandular myoepithelial cells (GMECs) have a unique capacity to specify ionocytes enriched in the Cftr channel. The Lef1 transcription factor was both necessary and sufficient to promote GMEC specification of ionocytes in mouse trachea, 3D organoids, and/or air-liquid interface (ALI) cultures. Overexpression of Lef1 (Lef1^OE^) in GMECs produced ionocyte-enriched epithelia with enhanced Cftr-mediated chloride, bicarbonate, and fluid transport properties. Moreover, *Foxi1* knockout in Lef1^OE^ GMECs prevented ionocyte formation while promoting specification of Cftr-expressing secretory cells. These findings provide a progenitor relationship for the anatomical association of ionocytes with SMGs and a transcriptional framework for coordinated specification of CFTR-expressing ionocytes and secretory cells.

**Highlights:** • Glandular myoepithelial cells give rise to pulmonary ionocytes in submucosal glands and the surface airway epithelium of mice.

• Disruption of *Lef1* in SMG progenitors decreases ionocyte specification and reduces CFTR-mediated anion transport.

• Overexpression of the Lef1 transcription factor in glandular progenitors promotes ionocyte frequency, CFTR-mediated fluid transport, and chloride and bicarbonate conductance.

•Disruption of *Foxi1* in SMG progenitors abolishes Lef1-dependent ionocyte specification and reduces CFTR-mediated anion transport.

## INTRODUCTION

Lineage tracing studies in mouse trachea have shown that myoepithelial cells in airway submucosal glands (GMECs) act as reserve stem cells that can specify into at least seven cell types that populate submucosal glands (ductal, mucous and serous acinar cells) and the surface airway epithelium (basal, club, goblet and ciliated cells) after severe airway injury ^1,2^. This process is mediated by the Wnt-regulated transcription factor lymphocyte enhancer binding factor 1 (Lef1) ^1^. Additionally, ectopic expression of Lef1 is sufficient to induce proliferation and specification of GMECs into various cell types without injury ^1^.

Pulmonary ionocytes are rare airway epithelial cells (0.5-1%) in proximal mouse trachea ^3^ and throughout cartilaginous human airways ^4^. Similar to intercalated cells in renal collecting ducts ^5^ and fish gill ionocytes ^6^, pulmonary ionocytes have a unique composition of ion channels suggestive of osmoregulatory function at the airway surface. Among these enriched channels is the cystic fibrosis transmembrane conductance regulator (Cftr) ^3,7^, in which mutations lead to cystic fibrosis ^8^. Thus, there is a particular interest in defining the origins of pulmonary ionocytes and their functional ion transport roles in regulating airway surface liquid composition and mucus rheology. Unlike renal collecting duct intercalated cells ^9,10^, the origins and pathways that regulate specification of pulmonary ionocyte are only partially understood. Previous studies have demonstrated that human, ferret, and mouse airway surface basal stem cells have the capacity to differentiate into pulmonary ionocytes in a Foxi1-dependent manner^3,7^.

Pulmonary ionocytes were first identified as CFTR-rich cells found to be enriched in airway submucosal glands (SMGs) ducts and the surrounding surface airway epithelium ^11^. The anatomical distribution of airway SMGs, and thus GMECs, is restricted to the proximal trachea in mice ^12,13^, whereas SMGs are found throughout the cartilaginous airways in humans ^13–15^, pigs ^13^ and ferrets ^16^. Despite species-specific differences in the abundance of SMGs, the association of pulmonary ionocytes with SMGs is conserved across species ^3,4^ and suggests the potential for a functional relationship between the anatomical location of ionocytes and SMGs. Notably, human pulmonary ionocytes harbor the highest expression of LEF1 among all airway epithelial cells ^7^. Given that Lef1 is also required for GMEC-mediated airway repair ^1,17^, we hypothesized that a Lef1-dependent lineage relationship between GMECs and pulmonary ionocytes exists.

In this work, we show that the Lef1 transcription factor is both necessary and sufficient to promote GMEC specification of ionocytes. Moreover, Lef1 overexpression (Lef1^OE^) in GMECs enhances ionocyte specification in mouse trachea, organoid culture, and polarized air-liquid interface (ALI) cultures while inhibiting the formation of Cftr-expressing secretory cells. Increased ionocyte abundance in Lef1^OE^ cultures enhanced their Cftr-mediated swelling response in organoid cultures and chloride and bicarbonate conductance in ALI cultures. Enhanced ionocyte specification and Cftr-mediated anion transport observed in Lef1^OE^ GMEC cultures was abolished by deletion of *Foxi1*, yet restored Cftr-expressing secretory cells to WT levels. These studies demonstrate a strong relationship between Cftr function and ionocyte presence in airway epithelia. Moreover, they provide a transcription framework by which Lef1 and Foxi1 coordinate specification of Cftr-expressing ionocytes and secretory cells by glandular progenitors.

## RESULTS

### Ectopic Lef1 expression in SMG progenitors promotes ionocyte specification and anion conductance

Wnt/Lef1 signaling plays a critical role in regulating the commitment of multipotent glandular progenitor cells to proliferate and differentiate into multiple airway cell types. However, the capacity to which glandular progenitors differentiate into pulmonary ionocytes remains unknown. Given that Lef1 is highly expressed in pulmonary ionocytes, we sought to evaluate the consequences of Lef1^OE^ on organoid formation and ionocyte specification by SMG progenitors. We prepared bulk SMG cells from eight-week-old *Lef1*-KI::*Rosa26*-LsL-nT mice and then treated them with tat-Cre recombinase and enriched by FACS for Tomato^+^ GFP^−^ (Lef1^OE^) cells. SMG progenitors from WT (Tomato^−^ GFP^+^) and Lef1^OE^ (Tomato^+^ GFP^−^) were then embedded at a 1:1 ratio (Lef1^OE^:WT) in Matrigel on transwell inserts. Differentiated Lef1^OE^ SMG progenitors (Tomato^+^) formed larger organoids with lumens compared to compact WT (GFP^+^) counterparts (Figure 1A–C). Stimulation of Cftr with the cAMP agonist forskolin resulted in swelling of Lef1^OE^ organoids but not WT organoids (Figure 1D–F). After seven weeks in culture, Lef1^OE^ organoids showed significantly a higher percentage of Foxi1^+^ ionocytes (Figure 1G–I) and had higher expression of ionocyte marker transcripts (*Foxi1*, *Bsnd* and *Cftr*) than WT counterparts (Figure 1J–L). These results demonstrate that Lef1 expression in bulk SMG progenitors promotes ionocyte specification and fluid transport properties in organoids typically associated with Cftr function.

**Figure 1:**
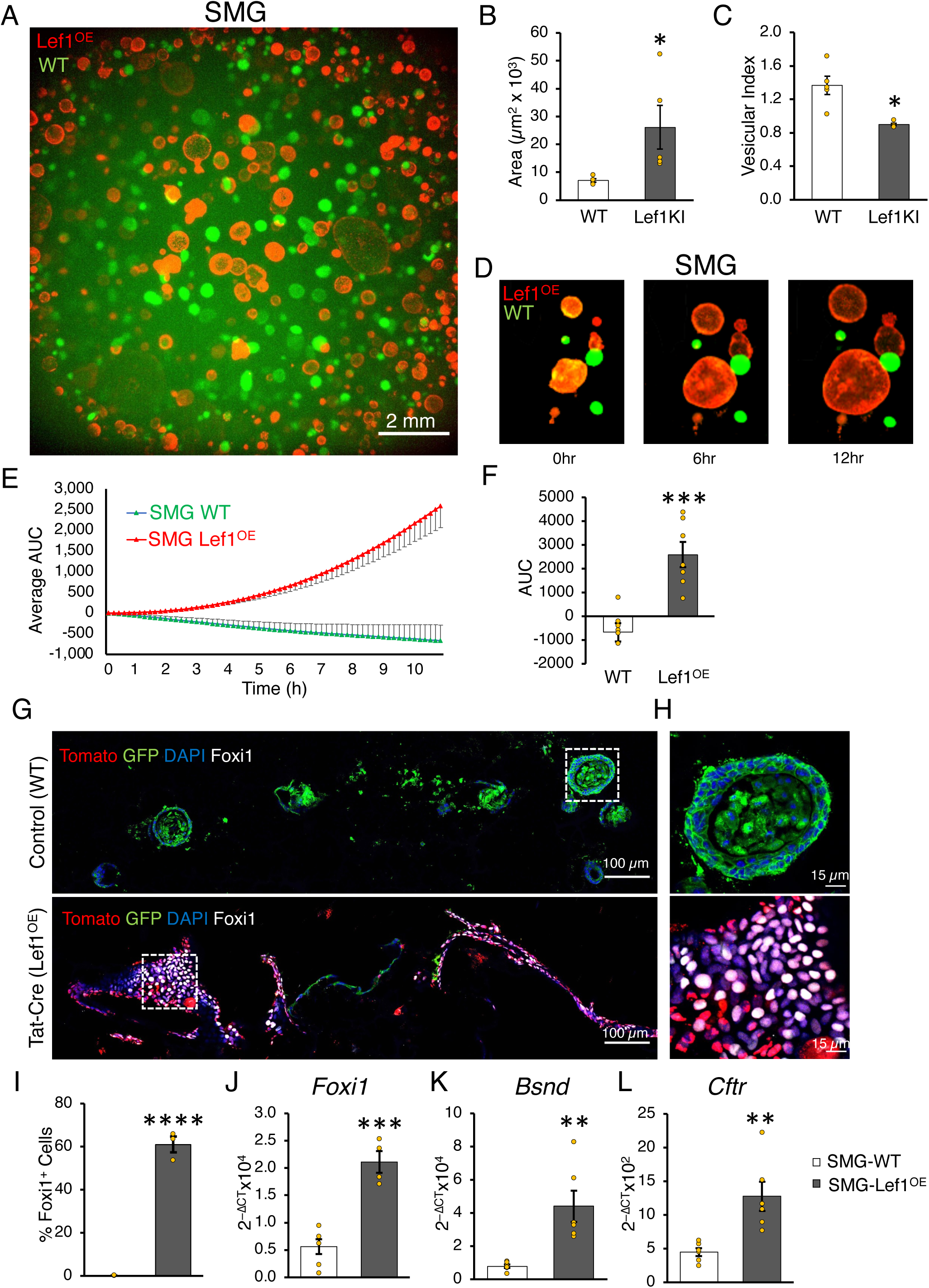
Lef1 overexpression in SMG progenitors promotes organoid formation and swelling. SMG progenitors were prepared from tracheas of eight-week-old *Lef1*-KI::*Rosa26*-LsL-nT mice. SMG progenitors were maintained in modified SAGM. SMG progenitors were treated with Tat-Cre to induce Lef1 overexpression (Lef1^OE^). Cells were sorted for Lef1^OE^ Tomato^+^ SMG progenitors. Wildtype (WT) EGFP^+^ cells were from the same cell preparation, but not treated with Tat-Cre. (**A–F**) Cells were plated at a 1:1 ratio (Lef1^OE^:WT) at a density of 3x10^4^ cells transwell^−1^ in matrigel. Transwells were grown in SAGM + supplements for one week, then switched to PneumaCult ALI for another two weeks. (**A**) A low power photomicrograph of a transwell of mixed SMG progenitors two weeks after differentiation. (**B**) Quantification of the average area of WT (green) and Lef1^OE^ (Red) organoids (n=5 donor groups). (**C**) Calculation of vesicular index (Intensity of the central part of organoid / Intensity of the organoid boundary) (n=5 donor groups). (**D**) Time series photomicrographs of organoid swelling over a 12-hour period after forskolin (10 µM) stimulation to cAMP-induce CFTR activity. (**E**) Swelling of SMG progenitor organoids in response to 10 µM forskolin for the indicated genotypes, measured as area under the curve (AUC) over a 12-hour period. (**F**) Quantification of AUC at the last time point (n=7 donor groups). (**G–L**) WT or Lef1^OE^ SMG progenitors were plated at a density of 3x10^4^ cells per transwell in matrigel. Transwells were grown in SAGM + supplements for one week and then maintained in PneumaCult ALI for another five weeks. Organoids were then processed as fresh frozen sections for immunofluorescence (**G–I**) or RNA isolation for RT-qPCR (**J–L**). (**G-I**) Immunofluorescence staining of organoid sections. (**G**) Photomicrograph of organoid sections stained for Tomato (Red), EGFP (Green) and FoxI1 (White). (**H**) A higher magnification of the boxed areas in G. (**I**) Quantification of Foxi1^+^ cells (n=3 donors). (**J–L**) qPCR of *Foxi1* (**J**), *Bsnd* (**K**) and *Cftr* (**L**) (n=6 donor groups). \**P*<0.05, \*\**P*<0.01, \*\*\**P*<0.001 and \*\*\*\**P*<0.0001, two-tailed Mann Whitney test. Error bars indicate SEM.

We then sought to evaluate the biologic properties of polarized Lef1^OE^ SMG progenitors that were differentiated at an air-liquid interface. Indeed, polarization and differentiation of Lef1^OE^ SMG progenitors at ALI produced more Foxi1^+^, Bsnd^+^ and Foxi1^+^ Bsnd^+^ cells (Figure S1A–D) and expressed higher levels of ionocyte markers (*Foxi1*, *Bsnd* and *Cftr*) (Figure S1E–G), compared to WT cultures. Lef1^OE^ SMG progenitor-derived ALI cultures also produced greater Cftr-dependent chloride and bicarbonate currents in response to cAMP agonists (IBMX/forskolin) than WT counterparts (Figure S1H–K). Therefore, Lef1 expression in SMG progenitors promoted ionocyte specification to produce epithelia with enhanced Cftr-dependent ion transport properties.

### Lef1 expression specifically in GMECs promotes ionocyte specification upon differentiation, giving rise to epithelia with enhanced anion conductance properties

Given that GMECs are thought to be the primary reserve stem cell population of SMGs and also utilize Lef1 during lineage commitment and multipotent repair of the mouse tracheal epithelium ^1^, we hypothesized that GMECs were the progenitor giving rise to pulmonary ionocytes. To this end, we utilized two Cre drivers (*Acta2*^CreERT2^ or *Myh11*^CreERT2^) to fate map GMECs on either *Lef1*-KI or *Rosa26*^mTmG^ backgrounds following tamoxifen injection. SMG progenitors were then prepared, and lineage traced Lef1^OE^ (GFP^−^) and WT (GFP^+^) GMECs were enriched by FACS. WT and Lef1^OE^ GMECs were then polarized in ALI cultures for seven weeks. Lef1^OE^ GMECs from both Cre drivers formed significantly more Foxi1^+^ Bsnd^−^, Bsnd^+^ Foxi1^−^ and Foxi1^+^ Bsnd^+^ ionocytes (Figure 2A–D) and expressed higher levels of ionocyte marker transcripts (*Bsnd*, *Foxi1* and *Cftr*) (Figure 2F–H,J-L) than WT counterparts. By contrast, expression of the α subunit of Enac (Scnn1a) was not different between groups (Figure 2E, I). Polarization and differentiation of Lef1^OE^ GMECs in ALI cultures gave rise to epithelia with significantly enhanced Cftr-dependent chloride (Figure S2A,B) and bicarbonate (Figure S2C,D) currents than the WT counterparts.

**Figure 2:**
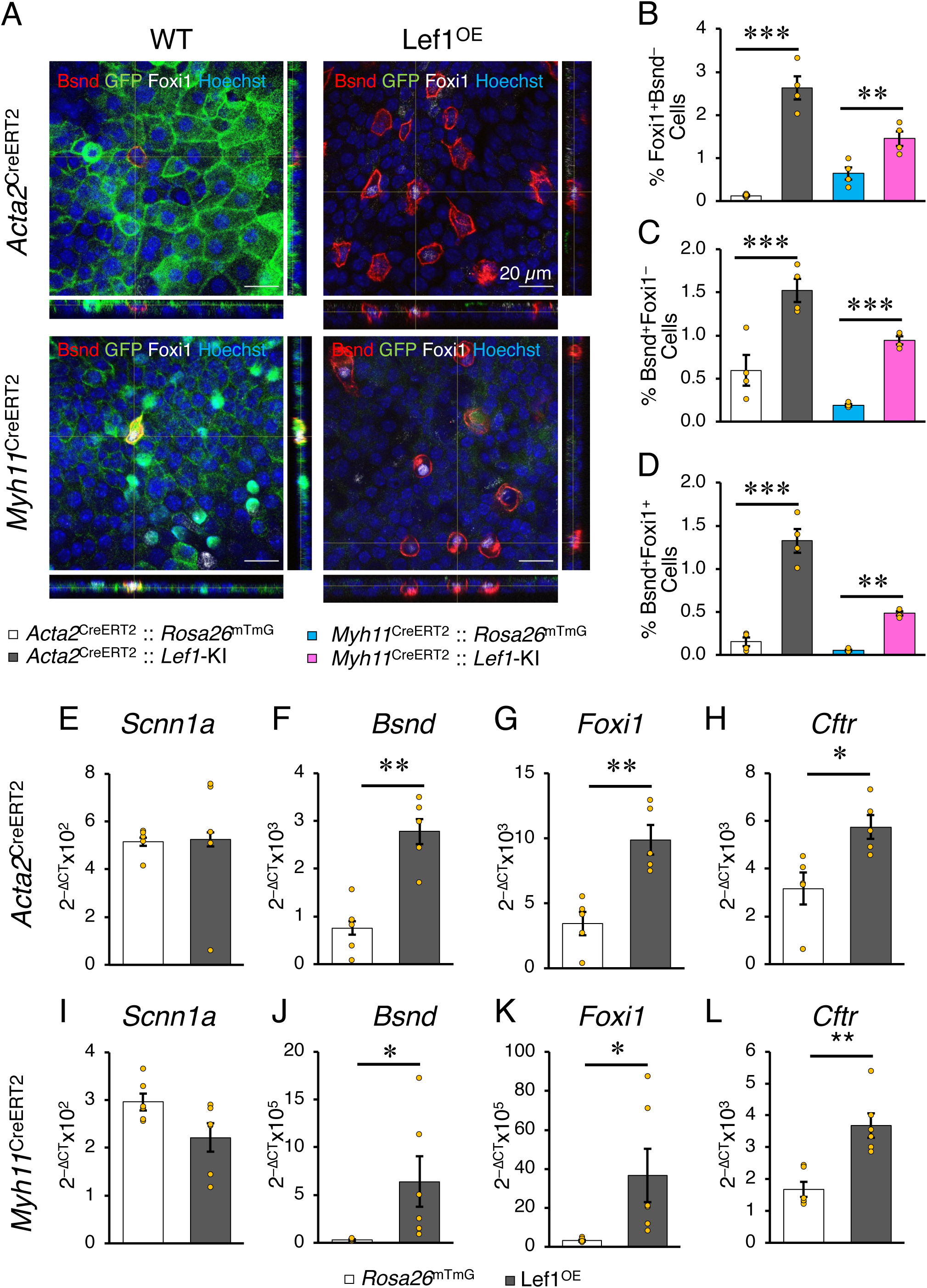
Lef1 overexpression in glandular myoepithelial cells (GMECs) promotes ionocyte specification. (**A-L**) *Acta2*^CreERT2^ or *Myh11*^CreERT2^ mice were bred to either *Lef1*-KI or *Rosa26*^mTmG^ mice. Double transgenic mice of all breeds received IP tamoxifen injection for 5 consecutive days. Ten days later all Mice were sacrificed and SMG progenitors were prepared and sorted to isolate GMECs of all genotypes. GMECs were plated on transwell membrane and ALI culture was established. Six weeks later ALI cultures underwent immunofluorescence staining or RNA preparation for RT-qPCR. (**A-D**). Photomicrographs with orthogonal views of whole mount ALI cultures immunolabelled for Bsnd (Red), GFP (Green), and Foxi1 (white) (**A**) and quantification of Foxi1^+^ Bsnd^−^ (**B**), Bsnd^+^ Foxi1^−^ (**C**) and Foxi1^+^ Bsnd^+^ (**D**) cells (n=4 donor groups). (**E-L**) RT-qPCR for *Scnn1a* (**E and I**), *Bsnd* (**F and J**), *Foxi1* (**G and K**) and *Cftr* (**H and L**) in *Acta2*^CreERT2^ (**E-H**) and *Myh11*^CreERT2^ (**I-L**) Cre driver genotypes (n=6 donor groups). \**P*<0.05, \*\**P*<0.01, \*\*\**P*<0.001 and \*\*\*\**P*<0.0001, two-way ANOVA followed by Bonferroni posttest (**B-D**) and two-tailed Mann Whitney test (**E-L**). Error bars indicate SEM.

To confirm *in vivo* that Lef1^OE^ GMECs differentiate into greater numbers of ionocytes than WT counterparts, we performed whole mount immunofluorescence staining of tracheas from *Acta2*^CreERT2^::*Lef1*-KI, *Myh11*^CreERT2^::*Lef1*-KI, *Acta2*^CreERT2^::*Rosa26*^mTmG^, and *Myh11*^CreERT2^::*Rosa26*^mTmG^ mice at 21 days after tamoxifen injection. Lef1^OE^ GMECs gave rise to a greater number of lineage-positive Bsnd- expressing ionocytes within SMGs and the surface airway epithelium, than did the WT counterparts (Figure 3A-C). By contrast, abundances of untraced Bsnd^+^ ionocytes were not significantly different between Lef1^OE^ and WT mice (Figure 3 D); these cells likely represent pre-existing ionocytes prior to tamoxifen treatment. Thus, Lef1 overexpression in GMECs promotes ionocyte specification *in vitro* and *in vivo*.

**Figure 3:**
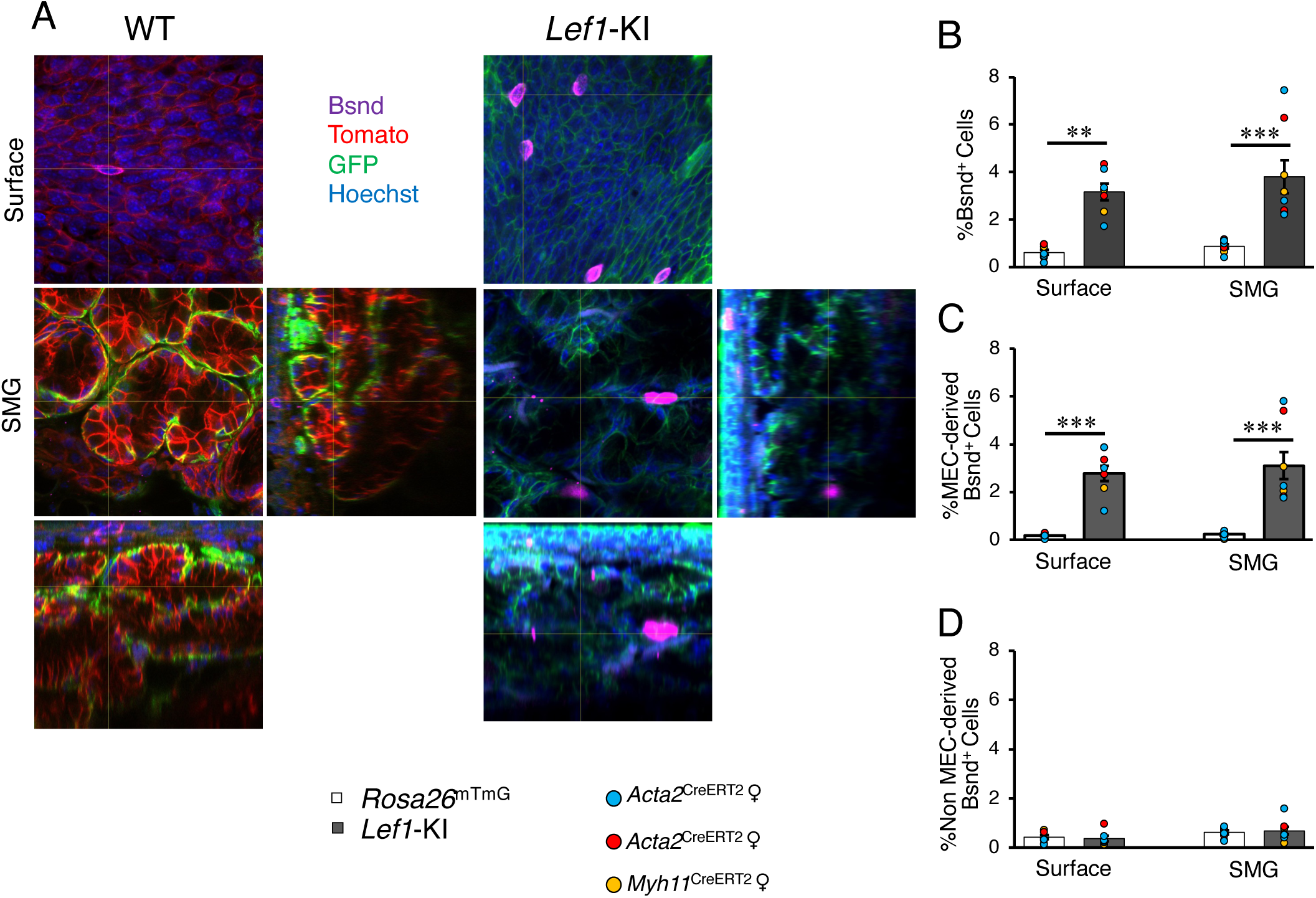
Lef1 overexpression in GMECs promotes ionocyte specification *in vivo*. (**A-D**) *Acta2*^CreERT2^ or *Myh11*^CreERT2^ mice were bred to either *Lef1*-KI or *Rosa26*^mTmG^ mice. Double transgenic mice of all breeds received IP tamoxifen injection for 5 consecutive days. Mice were sacrificed 21 days later and tracheas underwent whole mount immunofluorescence staining. (**A**) Orthogonal views of 3D images from whole mount tracheas immunolabelled for Bsnd (Magenta), EGFP (Green), and Tomato (Red). (**B-D**) Quantification of total Bsnd^+^ cells (**B**), Bsnd^+^ cells of GMEC origin (**C**), and Bsnd^+^ cells of non-GMEC origin (**D**) both on the airway surface and at the level of the submucosal gland (n=3, 2, and 2 donor groups of *Acta2*^CreERT2^ females “Blue circles”, males “Red circles” and *Myh11*^CreERT2^ males “Yellow circles”, respectively). Note: The *Myh11*^CreERT2^ locus is on the Y-chromosome so only males can be used for studies. \*\**P*<0.01, and \*\*\**P*<0.001, two-way ANOVA followed by Bonferroni posttest. Error bars indicate SEM.

### Foxi1 is required for Lef1-dependent ionocyte specification from SMG progenitors

Foxi1 is a transcription factor required for ionocyte differentiation from basal cells ^3^. We hypothesized that Lef1 functioned upstream of Foxi1 in the specification of ionocytes from GMECs. To test this hypothesis, we prepared SMG progenitors from *H11*^Cas9^::Lef1-KI double homozygous mice and used CRISPR/Cas9 to disrupt the *Foxi1* gene in Lef1^OE^ SMG progenitors. To facilitate simultaneous Lef1^OE^ and *Foxi1* disruption, we co-transfected SMG progenitors with sgRNAs targeting LoxP and *Foxi1* (herein called *Lef1*^OE^*Foxi1*^KO^ cells). Controls included SMG progenitors transfected with LoxP sgRNA alone (Lef1^OE^) or GFP sgRNA without (WT) or with (*Foxi1*^KO^) an sgRNA targeting the *Foxi1* locus. Analysis of *Foxi1* targeting efficiency using the inference of Cas9 editing (ICE) online tool confirmed *Foxi1* disruption in ∼90% of cells (Figure S3A–E). SMG progenitors of various genotypes were enriched for gene converted cells by FACS (EGFP*^−^*) and then polarized at an air-liquid interface.

Immunofluorescence staining of whole mount ALI culture membranes demonstrated that *Lef1*^OE^*Foxi1*^KO^ epithelia contained significantly (*P*<0.001) reduced numbers of Foxi1^+^Bsnd^+^ Cftr^+^, Bsnd^+^Cftr^+^, Foxi1^+^Cftr^+^, and Foxi1^+^Bsnd^+^ ionocytes compared to *Lef1*^OE^ epithelia (Figure 4A–E). While the magnitude of changes was smaller, *Foxi1*^KO^ epithelia also had reduced numbers of these cellular phenotypes compared to WT epithelia. Notably, the percentage of Cftr^+^ Bsnd^−^ Foxi1^−^ cells in *Lef1*^OE^ epithelia was significantly (*P*<0.05) reduced compared to WT and disruption of *Foxi1* in *Lef1*^OE^*Foxi1*^KO^ epithelia led to recovery (*P*<0.05) of Cftr^+^ Bsnd^−^ Foxi1^−^ cells to WT levels (Figure 4F). The opposite trend occurred in the absence of Lef1^OE^, where *Foxi1*^KO^ epithelia had reduced (*P*<0.05) numbers of Cftr^+^Bsnd^−^Foxi1^−^ cells compared to WT epithelia (Figure 4F). To characterize the Cftr^+^Bsnd^−^Foxi1^−^ cell type, ALI cultures were immunolabeled for secretoglobin family 1a member 1 (Scgb1a1), Muc5b, Cftr, and Bsnd. These studies showed that the majority of Cftr^+^Bsnd^−^Foxi1^−^ cells were Cftr^+^Scgb1a1^+^ club cells (Figure S3F–H). Disruption of *Foxi1* also significantly decreased expression of ionocyte marker transcripts (*Foxi1*, *Bsnd,* and *Cftr*) in ALI cultures derived from both WT and Lef1^OE^ SMG progenitors (Figure 4G–I). Finally, disruption of *Foxi1* resulted in an ∼50% reduction of Cftr-dependent chloride currents in both *Foxi1*^KO^ and *Lef1*^OE^*Foxi1*^KO^ epithelia as compared to their WT or *Lef1*^OE^ controls, respectively (Figure 4J–K). Thus, Lef1-dependent increases in ionocyte specification and Cftr function require Foxi1.

**Figure 4:**
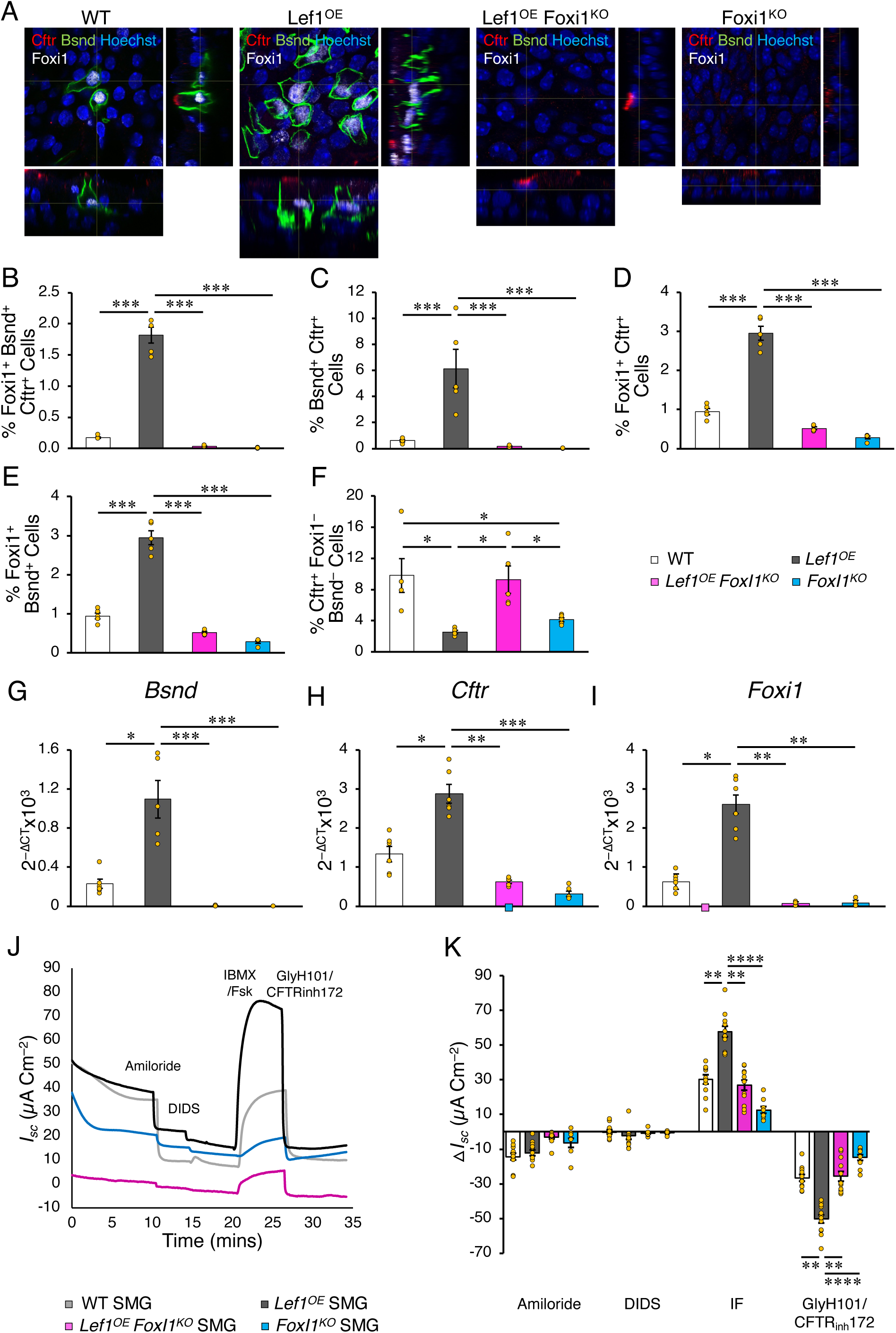
Lef1-mediated enhancement of CFTR-mediated anion conductance is ionocyte dependent. *H11*^Cas9^ mice were bred to homozygosity on the Lef1-KI background. Eight-week-old double transgenic mice were sacrificed, and SMG progenitors were prepared. SMG progenitors were transfected with CRISPR sgRNA targeting LoxP or GFP with ± CRISPR sgRNA targeting the *Foxi1* locus. The resultant SMG progenitors were *Lef1^OE^* (LoxP sgRNA alone), *Lef1^OE^ Foxi1^−/–^* (LoxP and *Foxi1* sgRNAs), *Foxi1^−/–^* (EGFP and *Foxi1* sgRNAs), or WT (EGFP sgRNA alone). SMG Progenitors of various genotypes were enriched gene converted cells by FACS (EGFP*^−^*) and then polarized at an air-liquid interface. (**A**) Orthogonal views of 3D images from whole mount ALI membranes immunolabelled for Cftr (Red), Bsnd (Green), and Foxi1 (White). (**B-F**) Quantification of Foxi1^+^ Bsnd^+^ Cftr^+^ (**B**), Bsnd^+^ Cftr^+^ (**C**), Foxi1^+^ Cftr^+^ (**D**), Foxi1^+^ Bsnd^+^ (**E**), and Cftr^+^ Foxi1^−^ Bsnd^−^ (**F**) cells (n=5 donor groups). (**G-I**) RT-qPCR for *Bsnd* (**G**), *Cftr* (**H**) and *Foxi1* (**I**) mRNA. (**J and K**) Short-circuit current (*I*_sc_) measurements under symmetrical chloride conditions performed on polarized SMG progenitors grown at an air-liquid interface. Shown are traces of representative experiments (**J**) and the changes in *I*_sc_ (Δ*I*_sc_) (**K**) in response to sequential addition of amiloride, 4,4ʹ- diisothiocyanatostilbene2,2ʹ-disulfonic acid (DIDS), forskolin + 3-isobutyl-1-methylxanthine (IBMX/Fsk), and GlyH101 + CFTR_Inh_ 172 (CFTR inhibitors), as indicated for the given genotypes (n= 11 donor groups). \*\*\**P*<0.001, \**P*<0.05, \*\**P*<0.01, and \*\*\*\**P*<0.0001, one-way ANOVA followed by Bonferroni posttest. Error bars indicate SEM.

Although Lef1^OE^ gain of function demonstrates Lef1 is sufficient to induce ionocytes from glandular progenitors, it remained unclear if Lef1 was necessary for ionocyte specification. To this end, we evaluated the capacity of *Lef1^−/–^* glandular progenitors to generate ionocytes using *Rosa26*^CreERT2^::*Rosa26*^mTmG^::*Lef1*- cKO triple transgenic mice. Proliferating glandular progenitors were plated at ALI in the presence of 4- hydroxytamoxifen for the duration of the experiment and compared to uninduced donor matched cells. Induction efficiency in the *Lef1^−/–^* cultures was confirmed by mT to mG conversion and remained at >98%. Indeed, *Lef1^−/–^* SMG progenitors differentiated into significantly (*P*<0.01) fewer ionocytes (Figure S4A–E) and had significantly (*P*<0.01) reduced Cftr-inducible chloride current (Figure S4F–G) than uninduced WT polarized ALI cultures. Thus, glandular progenitors require Lef1 to differentiate into ionocytes and Lef1 appears to act upstream to Foxi1 in the specification transcriptional network.

## DISCUSSION

Airway stem cells reside in multiple unique niches throughout the lung and play important regenerative roles in maintaining epithelial barrier function following injury ^18^. In the proximal conducting airways, multipotent basal cells are classically defined as the first responding stem cell compartment that facilitates repair after injury or infection ^19^. However, following severe injury and loss of surface airway basal cells, GMECs are mobilized to proliferate, migrate out of their niche, and specify into multiple cell types that populate the submucosal glands and the surface airway epithelium ^1,2^.

Pulmonary ionocytes and secretory cells are the predominant differentiated cell types in the human cartilaginous airways that express CFTR. These CFTR expressing cell types are thought to play a critical role in regulating airway surface liquid volume and mucus rheology required for proper mucociliary clearance (MCC) ^20^. SMGs also play very important roles in mucociliary clearance since they are the predominant site of mucus and liquid secretions into the airways ^20^. While these airway clearance mechanisms are notably disrupted in cystic fibrosis, they also share pathologic components with other forms of chronic obstructive pulmonary disease (COPD) ^21^.

Increasing interest in cell and gene therapies for cystic fibrosis have raised important questions about whether CFTR-expressing pulmonary ionocytes are appropriate targets for such therapies ^22^. For example, what are the mechanisms by which such rare cells function within a multi-cellular epithelium to regulate ion transport and fluid movement? Will proper regulation of CFTR within cell types that express this gene be required for effective gene therapy? If so, gene editing of ionocyte progenitors may be required to properly regulate CFTR functions. Here we show that GMECs are a progenitor cell population that gives rise to pulmonary ionocytes. Furthermore, we demonstrate that the Lef1 transcription factor is both required (Figure S4) and sufficient (Figure 1-4) for specifying pulmonary ionocytes from SMG progenitors. Given that Lef1^OE^ SMG progenitors require Foxi1 for the specification of ionocytes and Lef1 is also necessary for the specification event, Lef1 appears to function upstream to Foxi1 in the cell fate decision by GMECs. Since GMECs acquire a basal stem cell-like phenotype in culture ^1^, we propose the GMECs may commit to forming a specialized progenitor that is uniquely suited to differentiate into ionocytes with specialized CFTR functions.

Our studies show that Lef1^OE^ SMG progenitors give rise to a higher percentage of ionocytes (Foxi1^+^) in organoid culture (∼60%) (Figure 1I) compared to ALI cultures derived from the same cells (∼13%) (Figure S1B and C). This might be due to the effect of the surrounding extracellular matrix (ECM) and/or the 3D organization of the SMG progenitors in culture. Our *in vivo* lineage-tracing experiments show that ∼30% (WT) or ∼82% (Lef1^OE^) of Bsnd^+^ ionocytes within tracheal submucosal glands originate from GMECs during a 21-day chase period at steady state (Figure 3C). Similarly, in the surface airway epithelium, the percentages of GMEC-lineage Bsnd^+^ ionocytes were 26% (WT) and 88% (Lef1^OE^) after the 21-day chase period. This relatively high level of GMEC to ionocyte conversion in WT trachea at homeostasis suggests a high rate of turnover of ionocytes, as compared to the estimated 6-month half-life of mouse tracheal ciliated cells ^23^. Given that ionocytes are enriched in mitochondria, presumably to meet the energy requirements for regulated ATP- and cAMP-dependent channels, the apparently high rate of turnover may be due to redox-dependent mitochondrial damage. These findings emphasize that GMECs may play a more active role in homeostatic specification of surface airway ionocytes than previously appreciated.

Lef1 is known to control the cell cycle of airway stem cells ^24^ and to promote proliferation and differentiation of GMECs ^1^. Additionally, single cell RNA sequencing studies have shown that LEF1 is enriched in human pulmonary ionocytes ^7^. Consistent with previous studies ^3^, knocking out *Foxi1* resulted in inhibition of ionocyte specification and decreased Cftr-mediated chloride conductance in both WT and Lef1^OE^ polarized SMG ALI cultures (Figure 4). Furthermore, based on Cftr, Bsnd, and Foxi1 co-expression, multiple ionocyte sub-populations appear to be downregulated in the absence of *Foxi1* (Figure 4B-E). Notably, Lef1^OE^ ALI cultures contained fewer Cftr**^+^**Scgb1a1**^+^**BSND**^−^** club cells, but disruption of *Foxi1* in Lef1^OE^ ALI cultures restored Cftr**^+^**Scgb1a1**^+^**BSND**^−^** club cells to WT levels (Figure S3F-H). In light of the fact that ALI cultures derived from Lef1 knockout mice specify large numbers of Scgb1a1^+^ and Muc5B^+^ secretory cells ^24^, our findings implicate Lef1 and Foxi1 in a transcriptional axis that governs specification of the two major Cftr-expressing cell types in the proximal airway (i.e., ionocytes and secretory cells). Although Lef1^OE^ Foxi1^KO^ ALI cultures showed a higher percentage of Cftr**^+^**Bsnd^−^Foxi1^−^ cells than Foxi1^KO^ cultures (Figure 4F), there was no significant difference in *Cftr* mRNA level or Cftr-mediated *I_sc_* between the two groups and *Cftr* mRNA expression generally correlated with changes in Cftr-mediated current across all four genotypes (Figure 4H,J,K). Thus, the difference in the abundance of Cftr**^+^**Scgb1a1**^+^** club cells between Lef1^OE^ Foxi1^KO^ and Foxi1^KO^ cultures had little impact on transepithelial currents, despite the fact that Cftr appeared to localize to the apical domain of club cells in both WT and Lef1^OE^ Foxi1^KO^ cultures (Figure S3F). These findings raise important questions about distinct functions of Cftr in ionocytes vs secretory cells.

A limitation of the current study is that mice served as the model species, and they do not recapitulate the pulmonary phenotype of CF in the absence of Cftr. However, the biology of GMECs ^1,2^ and Cftr-mediated SMGs secretion ^25–27^ appear conserved across multiple species, including mice, despite the fact that SMGs are only found in the proximal trachea of mice. Thus, the glandular progenitor that specifies ionocytes in the airway may also be conserved across species.

Our study provides new insights into the origins of SMG progenitors that seed pulmonary ionocytes in both SMGs and the surface airway epithelium and the role of Lef1 in this process. Thus, GMECs may be suitable targets for gene and stem cell therapy and manipulation of the Lef1 transcription factor to promote pulmonary ionocyte specification may have therapeutic application for improving mucociliary clearance in COPD patients and CF patients on CFTR modulators.

## ACKNOWLEDGMENTS

This work was supported by NIH grant (R01 HL165404, P01 HL152960, P30 DK054759 to J.F.E.); CFF grant (ENGELH20XX2 to JFE and CY) the Carver Chair in Molecular Medicine (to J.F.E.)

## AUTHOR CONTRIBUTIONS

Conceptualization, W.S., C.Y., J.F.E.; Methodology, W.S., C.Y., and J.F.E.; Analyses, W.S., C.Y., O.S., V.I., T.L., and F.Y.; Investigation, W.S., J.H., O.S., and G.F.; Resources, C.Y.; Writing – Original Draft, W.S., J.F.E.; Writing – Review & Editing, W.S., J.H., C.Y., and J.F.E.; Visualization, W.S., and J.F.E.; Supervision, J.F.E., and C.Y.; Project Administration, W.S., C.Y., and J.F.E.

## STAR * METHODS

### KEY RESOURCES TABLE

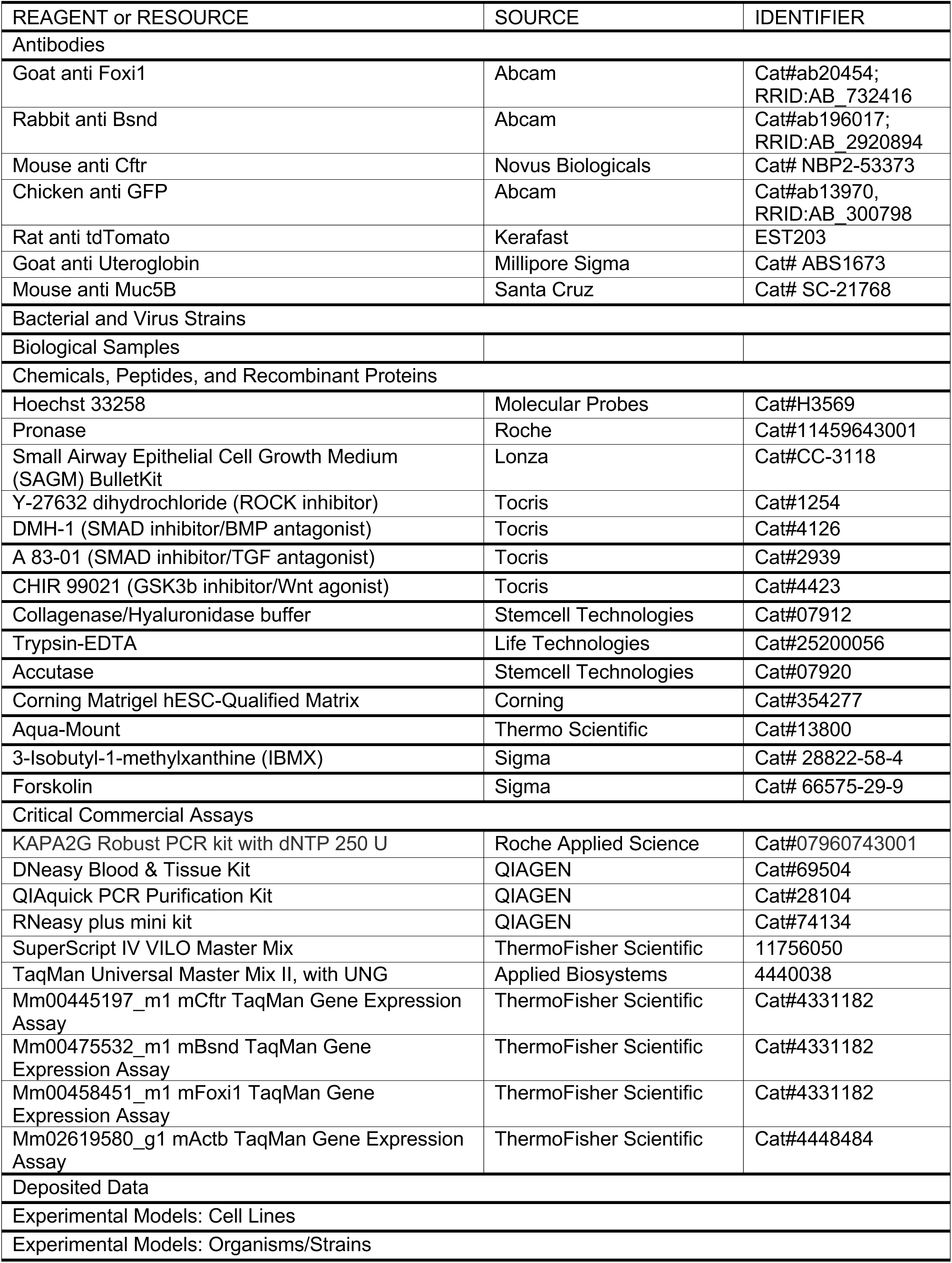
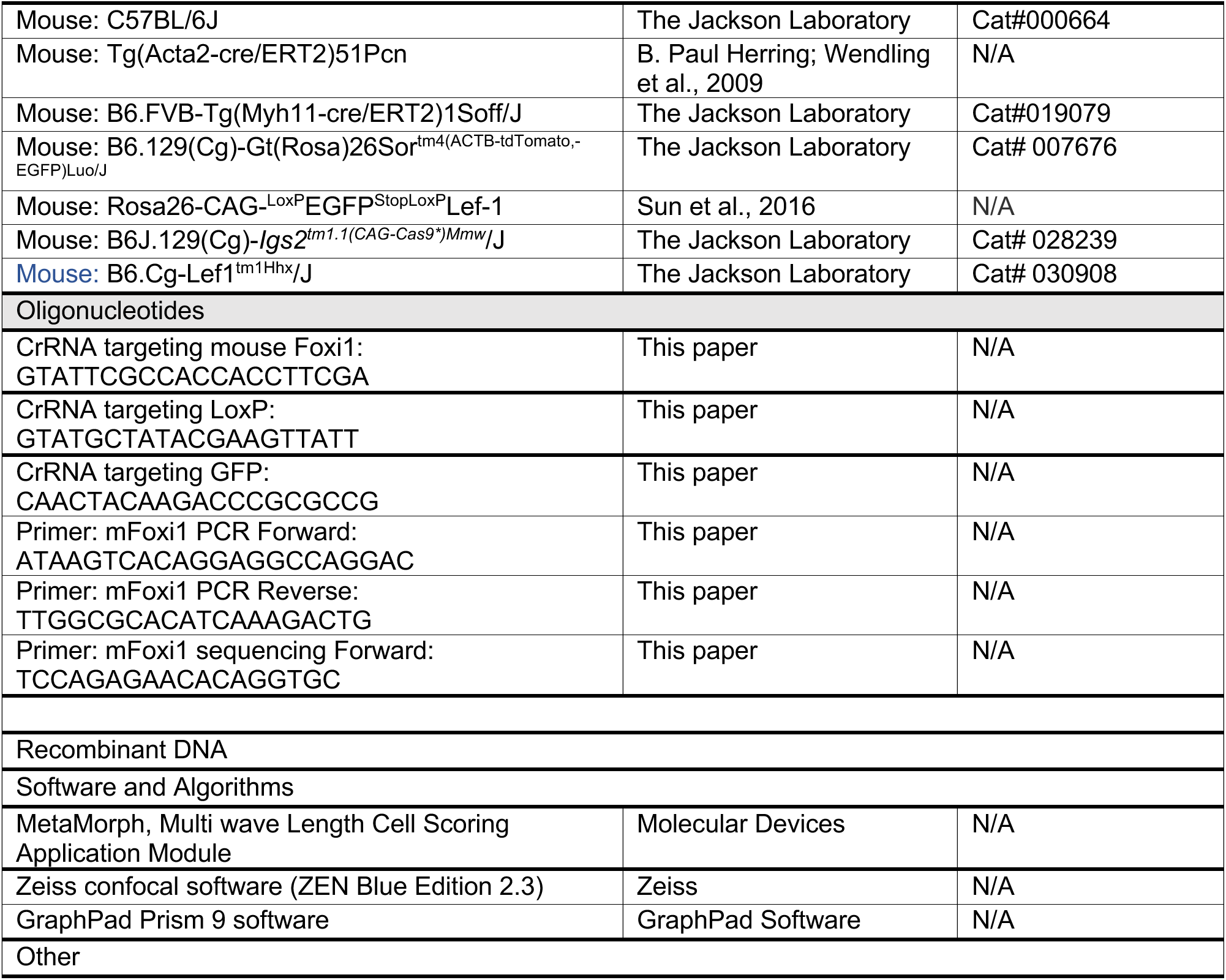

### RESOURCE AVAILABILITY

#### Lead contact

Further information and requests for resources and reagents should be directed to and will be fulfilled by the lead contacts, John F. Engelhardt and Charles Yeaman.

### EXPERIMENTAL MODEL AND SUBJECT DETAILS

#### Animal Studies

All procedures involving animals were performed in compliance with a protocol approved by the Institutional Animal Care and Use Committee of the University of Iowa and under institutional assurances including: AAALACi accreditation (#000833, since November 1994), USDA research facility registration (USDA No. 42-R-0004), and PHS Animal Welfare Assurance approval (D16-00009, A3021-01). Animals were maintained on a 12 h light/dark cycle and provided with food and water *ad libitum* in individually ventilated units in the specific-pathogen free facilities at the University of Iowa. Female and male (6 to 8 week-old) mice of the following genotypes and strains were used: B6.129(Cg)-Gt(Rosa)26Sor^tm4(ACTB-tdTomato,- EGFP)Luo^/J (*Rosa26*^mTmG^) Cre-reporter mice (The Jackson Laboratory, stock number 007676), *Myh11*^CreERT2^ (The Jackson Laboratory, stock number 019079) , B6.Cg-Lef1^tm1Hhx^/J (Lef1-cKO, The Jaxon Laboratory, stock number 030908), B6J.129(Cg)-*Igs2^tm1.1(CAG-Cas9*)Mmw^*/J (*H11*^Cas9^, The Jackson Laboratory, stock number 028239). For the purpose of this manuscript, this line is called, *Acta2*^CreERT2^ (generously provided by Dr. B. Paul Herring’s lab and maintained on a C57BL/6 background), and Gt(Rosa)26Sor^tm1(CAG-LEF1,-GFP*)Adj^ mice (maintained on a C57BL/6 background). *Acta2*-Cre^ERT2^ mice were described previously ^28^. Gt(Rosa)26Sor^tm1(CAG-LEF1,-GFP*)Adj^ knock-in mice were used to overexpress the human *LEF1* transgene in response to Cre and were described previously ^29^. For the purposes of this manuscript, this line is called Lef1-KI. In all studies both male and female mice were utilized with the exception that only male *Myh11*^CreERT2^::*Rosa26*^mTmG^ mice were evaluated, since the *Myh11^CreERT2^* transgene is on the Y chromosome. Animals were genotyped at weaning from tail clippings and identified using earhole punches. Experiments were initiated using mice that had no known health/immune concerns, were not involved in previous procedures, and were otherwise drug/test naive. For lineage tracing experiments Cre-mediated recombination was induced in mice by i.p. injection of 75 μg tamoxifen per gram bodyweight every 24 hr for a total of 5 consecutive days. Mice were allowed to recover for 5 or 21 days between tamoxifen treatment and sacrifice. Mice were induced with tamoxifen between 6-8 weeks of age.

#### Culture conditions for *in vitro* systems

Primary cells were cultured in Small Airway Epithelial Cell Growth Medium (SAGM) (Lonza, Cat#CC-3118) modified with the addition of 10 μM Y-27632 (Tocris, Cat#1254), 1 μM DMH-1 (Tocris, Cat#4126), 1 μM A83-01 (Tocris, Cat#2939), and 1 μM CHIR 99021 (Tocris, Cat#4423) on tissue culture plastic pre-treated with filter-sterilized laminin-enriched 804G-conditioned media as previously described ^30^.

### Method Details

#### Tissue Processing and Cell Isolation

Epithelia from resected mouse tracheae were isolated using a sequential enzymatic digestion strategy as previously described with slight modifications ^31^. Tracheae were opened longitudinally to expose the SAE and then digested in 1.5 mg/ml Pronase (Roche) in DMEM:F12 at 37°C for 60 minutes with gentle nutation. Tissues were gently agitated to remove SAE and then passed through a 100 μm cell strainer. The flow through containing SAE cells was changed into DMEM:F12 and then into modified SAGM (Lonza) ^30^ prior to plating for culture. The remaining tracheal tissue was then dissected with fine tip surgical scissors into tissue pieces ≤ 3 mm^3^ and further digested to isolate SMG cells after washing tissue fragments to remove lightly adherent cells (5 changes of DMEM:F12 by pipetting up and down using a 5 mL plastic pipette). Tissue fragments were then incubated in 2X Collagenase/Hyaluronidase buffer (StemCell Technologies, Inc.) diluted in DMEM:F12 at 37°C for 45 minutes with gentle nutation. Pre-warmed 0.25% Trypsin-EDTA (Life Technologies) was then added to the cell mixture to a final concentration of 0.05% Trypsin-EDTA and incubated for an additional 30 minutes at 37°C with gentle nutation. After pipetting up and down using a P1000 pipette, a single cell suspension was obtained by passing the cell mixture through a 100 μm cell strainer. The flow through containing SMG cells was changed into DMEM:F12 and then into modified small airway growth media (SAGM, Lonza) as indicated above prior to plating for culture. All centrifugations were performed at 250 × g for 7 min.

#### Tat-Cre treatment

SMG progenitors from *Lef1*-KI::*Rosa26*-LsL-nT mice were incubated with 5 µM Tat-Cre (Millipore Sigma) for three hours at 37°C, SMG progenitors were then washed twice with 1x PBS and watched daily for color change from GFP**^+^** to nTomato**^+^**. SMG cells were then sorted for Lef^OE^ nTom**^+^** using flow cytometric analysis.

#### Flow Cytometry

Flow cytometric analysis was performed on cultured primary SMG cells isolated from *Lef1*-KI::*Rosa26*-LsL- nT, *Acta2*^CreERT2^::*Rosa26*^mTmG^ or *Acta2*^CreERT2^::*Lef1*-KI mice. Cells were dissociated from plastic plates using Accutase (StemCell Technologies, Inc.), changed into HBSS containing 2% FBS, and passed through a 40 μm cell strainer. GFP**^+^** and Tomato**^+^** (*Acta2*^CreERT2^::*Rosa26*^mTmG^ mice) or GFP**^+^** and GFP**^−^** (*Acta2*^CreERT2^::*Lef1*-KI mice) cell populations were identified after gating for viability using Hoechst 33258 (Molecular Probes) at a final concentration of 4 μg/ml. Cells were analyzed and sorted on a FACS Aria II (BD Biosciences).

#### 3D SMG organoid culture

WT or Lef1^OE^ SMG progenitors were seeded at 3 x 10^4^ cells/0.33 cm^2^ transwell in 100 µl of modified SAGM and Matrigel (Corning) at 1:1 ratio. SMG progenitors were maintained in SAGM for seven days. After which, cells were differentiated in PneumaCult ALI (StemCell Technologies, Inc.) for another three or six weeks.

#### Organoid swelling assays

WT and Lef1^OE^ SMG progenitors were seeded at 1:1 ratio in Matrigel. Transwells were grown in SAGM + supplements for one week, then switched to PneumaCult ALI for another three weeks. After which, Matrigel was disrupted, and organoids were seeded into a 96-well plate. Forskolin-induced swelling (FIS) of WT and Lef1^OE^ organoids was assessed by measuring the response to forskolin (5 μM final concentration) overnight with images captured at 10-minute intervals (total 97 time points). FIS of organoids is expressed as the sum of the absolute area under the curve (AUC). Zeiss confocal software (ZEN Blue Edition 2.3) was utilized to calculate area (μm^2^) as previously described ^32^.

#### Immunofluorescence

Transwells of organoid cultures were snap frozen on dry ice after seven weeks in culture. Organoids in Matrigel were physically separated from the transwells while still frozen and embedded in OCT frozen blocks. Frozen sections were cut at 7 µm and mounted on slides. Frozen sections were fixed in 4% PFA in PBS for 30 minutes at room temperature prior to washing in three changes of PBS. Sections were boiled at 98°C for 35 minutes in a solution containing 10 mM sodium citrate, 0.2% tween 20 pH 6 for antigen retrieval (note that this antigen retrieval leads to a more diffuse GFP staining pattern in Lef1-KI organoids but is required to detect nuclear Foxi1). Slides were incubated in blocking buffer containing 20% normal donkey serum, 0.3% Triton X-100, and 1 mM CaCl_2_ in PBS for two hours at room temperature. The slides were incubated with a mixture of primary antibodies in diluent buffer containing 1% normal donkey serum, 0.3% Triton X-100, and 1 mM CaCl_2_ in PBS overnight at 4°C. Slides were washed in three changes of PBS and incubated with a mixture of secondary antibodies in diluent buffer for two hours at room temperature. Fluorescent images were collected with a Zeiss LSM 700 line-scanning confocal microscope (Carl Zeiss, Germany). Nuclei were stained using Hoechst 33342 (Invitrogen). Slides were mounted with Aqua-Mount (Thermo Scientific).

#### Air-liquid Interface Cultures

Expanded primary WT (GFP**^+^**) or FACS enriched Lef1^OE^ (nTomato**^+^**) SMG progenitor cells were grown at an air-liquid interface (ALI) on 0.33 cm^2^ polyester transwell membranes (Corning) that were pre-treated with 804G-conditioned media. Each well was seeded with 2 × 10^5^ cells suspended in modified SAGM media (see above). At 24 hr post-seeding, cultures were moved to an air-liquid interface and maintained with PneumaCult ALI media (StemCell Technologies, Inc.) for at least six weeks. For GMEC experiments, ALI cultures were also established using FACS purified WT GMEC (GFP**^+^** cells isolated from *Acta2*^CreERT2^::*Rosa26*^mTmG^ mice) and Lef1^OE^ MEC (GFP**^−^** cells isolated from *Acta2*^CreERT2^::*Lef1*-KI mice) or (GFP**^+^** cells isolated from *Myh11*^CreERT2^::*Rosa26*^mTmG^ mice) and Lef1^OE^ MEC (GFP**^−^** cells isolated from *Myh11*^CreERT2^::*Lef1*-KI mice).

#### Short-circuit current (*Isc*)measurements

Transepithelial short-circuit currents (*I_sc_*) were measured using a self-contained Ussing chamber system (Warner Instruments, CT) as previously described ^33^. Throughout the experiment the chamber was kept at 37 °C. For chloride conductance, epithelia were bathed on both surfaces with a solution containing 135 mM NaCl, 2.4 mM K_2_HPO_4_, 0.6 mM KH_2_PO_4_, 1.2 mM CaCl_2_, 1.2 mM MgCl_2_, 10 mM dextrose, and 5 mM Hepes (pH 7.4) and bubbled with air. For bicarbonate conductance, epithelia were a solution containing 118.9 mM sodium gluconate, 25 mM NaHCO_3_, 5 mM dextrose, 1 mM magnesium gluconate, 5 mM calcium gluconate, 0.6 mM KH_2_PO_4_, 2.4 mM K_2_HPO_4_ and bubbled with CO_2_. *I_sc_* was measured while amiloride (100 µM), DIDS (100 µM), IBMX (100 µM) plus Forskolin (10 µM), and GlyH101 (50 µM) plus CFTR_Inh_ 172 (50 µM) were added sequentially to the apical chamber.

#### Crispr/Cas9 Knockout

Crispr/Cas9 knockouts were performed in primary SMG progenitors isolated from *H11*^Cas9^::*Lef1*-KI mice. Cells were grown to 60-80% confluence in 6-well plates and transfected using 0.3 nMoles of gRNA against the LoxP or GFP and 0.6 nMoles of gRNA against the gene of interest (*Foxi1*). Transfection was done using 10µl of Lipofectamine RNAiMax (Thermofisher scientific). FACS was performed 10 days after transfection. DNA was isolated from sorted cells using DNeasy Blood and Tissue Kit (Qiagen) and the gene locus was amplified by PCR. PCR product was Sanger sequenced with a subsequent peak de-convolution using ICE analysis (Synthego).

#### Immunofluorescence of whole mount mouse tracheae

Mouse tracheae were cut open longitudinally and fixed in 4% PFA in PBS for 48 hours prior to washing in in three changes of PBS. Tracheae were incubated in blocking buffer containing 20% normal donkey serum, 0.3% Triton X-100, and 1 mM CaCl_2_ in PBS over night at 37°C with rocking. The tracheae were incubated with primary antibody (or a mixture of primary antibodies) in diluent buffer containing 1% normal donkey serum, 0.3% Triton X-100, and 1 mM CaCl_2_ in PBS overnight at 37°C. Tracheae were washed in three changes of PBS and incubated with a mixture of secondary antibodies in diluent buffer overnight at 37°C. Fluorescent images were collected with a Zeiss LSM 980 line-scanning confocal microscope (Carl Zeiss, Germany). Nuclei were stained using Hoechst 33342 (Invitrogen). Tracheae were mounted on a slide with Aqua-Mount (Thermo Scientific) and flattened between two magnets.

#### Immunofluorescence of whole mount ALI transwell membrane

Whole mount ALI cultures were fixed in 4% PFA in PBS for 48 hours prior to washing in three changes of PBS. Transwells were boiled at 98°C for 35 minutes in a solution containing 10 mM sodium citrate, 0.2% tween 20 pH 6 for antigen retrieval (note that this antigen retrieval leads to a more diffuse GFP staining pattern in Lef1-KI mice but is required to detect nuclear Foxi1). Transwells were incubated in blocking buffer containing 20% normal donkey serum, 0.3% Triton X-100, and 1 mM CaCl_2_ in PBS for two hours at room temperature. The transwells were incubated with a mixture of primary antibodies in diluent buffer containing 1% normal donkey serum, 0.3% Triton X-100, and 1 mM CaCl_2_ in PBS overnight at room temperature. Transwells were washed in three changes of PBS and incubated with a mixture of secondary antibodies in diluent buffer for two hours at room temperature. Fluorescent images were collected with a Zeiss LSM 700 line-scanning confocal microscope (Carl Zeiss, Germany). Nuclei were stained using Hoechst 33342 (Invitrogen). Membranes were mounted on a slide with Aqua-Mount (Thermo Scientific) and flattened between two magnets.

#### Reverse transcription and quantitative polymerase chain reactions (rtPCR and qPCR)

After six weeks in Pneumacult ALI medium, ALI or organoid cultures were snap frozen on dry ice and kept in RNA protect (Qiagen) at 4°C. Total RNA was prepared using the RNeasy Plus Mini Kit (Qiagen). cDNA was prepared from the total RNA using the SuperScript IV VILO Master Mix (ThermoFisher Scientific) according to the manufacturer’s protocol. Quantification of transcription levels relative to β-Actin (inner control) was performed using TaqMan Universal Master Mix II, with UNG (Applied Biosystems), according to the manufacturer’s protocol and utilizing the following gene expression assays: Mm02619580_g1 Actb 4448484 TaqMan Gene Expression Assay, SM VIC PL for *β-Actin*, Mm00458451_m1 Foxi1 4331182 TaqMan Gene Expression Assay, SM for *Foxi1*, Mm00445197_m1 Cftr 4331182 TaqMan Gene Expression Assay, SM for *Cftr* and Mm00475532_m1 Bsnd 4331182 TaqMan Gene Expression Assay, SM for *Bsnd* (ThermoFisher Scientific).

#### Assessment of ionocyte specification after disruption of *Lef1*

SMG progenitors were prepared from *Rosa26*^CreERT2^::*Rosa26*^mTmG^::*Lef1*-cKO triple transgenic mice and were plated on transwell inserts in SAGM + supplements for 3 days then maintained in PnumaCult ALI on the basolateral surface to establish air liquid interface (ALI) culture. Half of the transwells from each donor were maintained in medium containing 2 µg/ml 4-hydroxytamoxifen for the entire length of the experiment. Six weeks later, transwells of all groups underwent short circuit current measurement to assess anion conductance followed by fixation and wholemount immunofluorescence staining.

