## Supplementary Materials for "Lef1 Promotes Foxi1-Dependent Ionocyte Specification by Airway Submucosal Gland Progenitors"

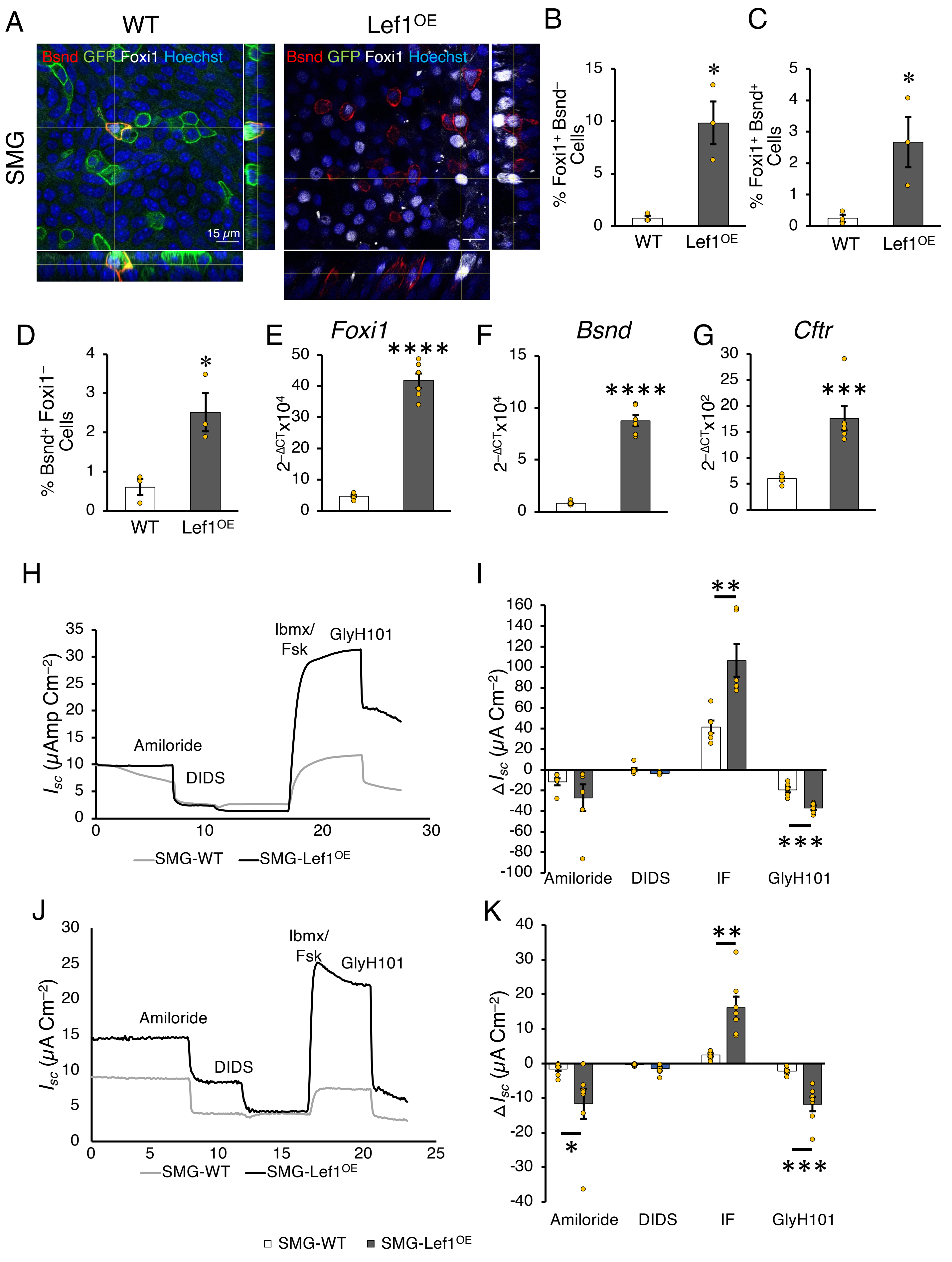
A
WT
Lef1OE
SMG
Bsnd GFP Foxi1 Hoechst
15 μm
B
% Foxi1+ Bsnd− Cells
C
% Foxi1+ Bsnd+ Cells
D
% Bsnd+ Foxi1− Cells
E
Foxi1
F
Bsnd
G
Cftr
H
Amiloride
DIDS
Ibmx/ Fsk
GlyH101
SMG-WT
SMG-Lef1OE
I
Amiloride
DIDS
IF
GlyH101
J
K
SMG-WT
SMG-Lef1OE

**Figure S1: Lef1 overexpression in SMG progenitors promotes ionocyte specification and CFTR-mediated anion conductance. Related to Figure 1.** (**A-K**) WT or Lef1$^{OE}$ SMG progenitors from the donors described in Fig. 1 were plated on transwell inserts in SAGM + supplements for 3 days then maintained in PnumaCult ALI on the basolateral surface to establish air liquid interface (ALI) culture. Six weeks later a set of tanswells underwent short circuit current measurement to assess anion conductance followed by fixation and wholemount immunofluorescence staining. Another set of transwells were used to prepare RNA for RT-qPCR. (**A–D**) Orthogonal views of 3D images of whole mount membranes stained for Foxi1 (White), Bsnd (Red) and EGFP (Green) (**A**) and quantification of Foxi1$^+$ Bsnd$^-$ (**B**), Foxi1$^+$ Bsnd$^+$ (**C**) and Foxi1$^-$ Bsnd$^+$ (**D**) cells (n=3 donor groups). (**E–G**) RT-qPCR for *Foxi1* (**E**), *Bsnd* (**F**) and *Cftr* (**G**) mRNA from ALI cultures (n=6 donor groups). (**H–K**) Short-circuit current ($I_{sc}$) measurements on polarized SMG progenitors grown at an air-liquid interface performed under symmetrical chloride (**H**) or symmetrical bicarbonate (**J**) conditions. Shown are traces of representative experiments (**H and J**) and the changes in $I_{sc}$ ($\Delta I_{sc}$) (**I and K**) in response to sequential addition of amiloride, 4,4′-diisothiocyanatostilbene2,2′-disulfonic acid (DIDS), forskolin + 3-isobutyl-1-methylxanthine (IBMX/FSK), and GlyH101, as indicated for the given genotypes. n= 5 and 7 donor groups in I and K respectively. *$P$<0.05, **$P$<0.01, ***$P$<0.001 and ****$P$<0.0001, two-tailed Mann Whitney test (**B-G**) and two-way ANOVA followed by Bonferroni posttest (**I and K**). Error bars indicate SEM.

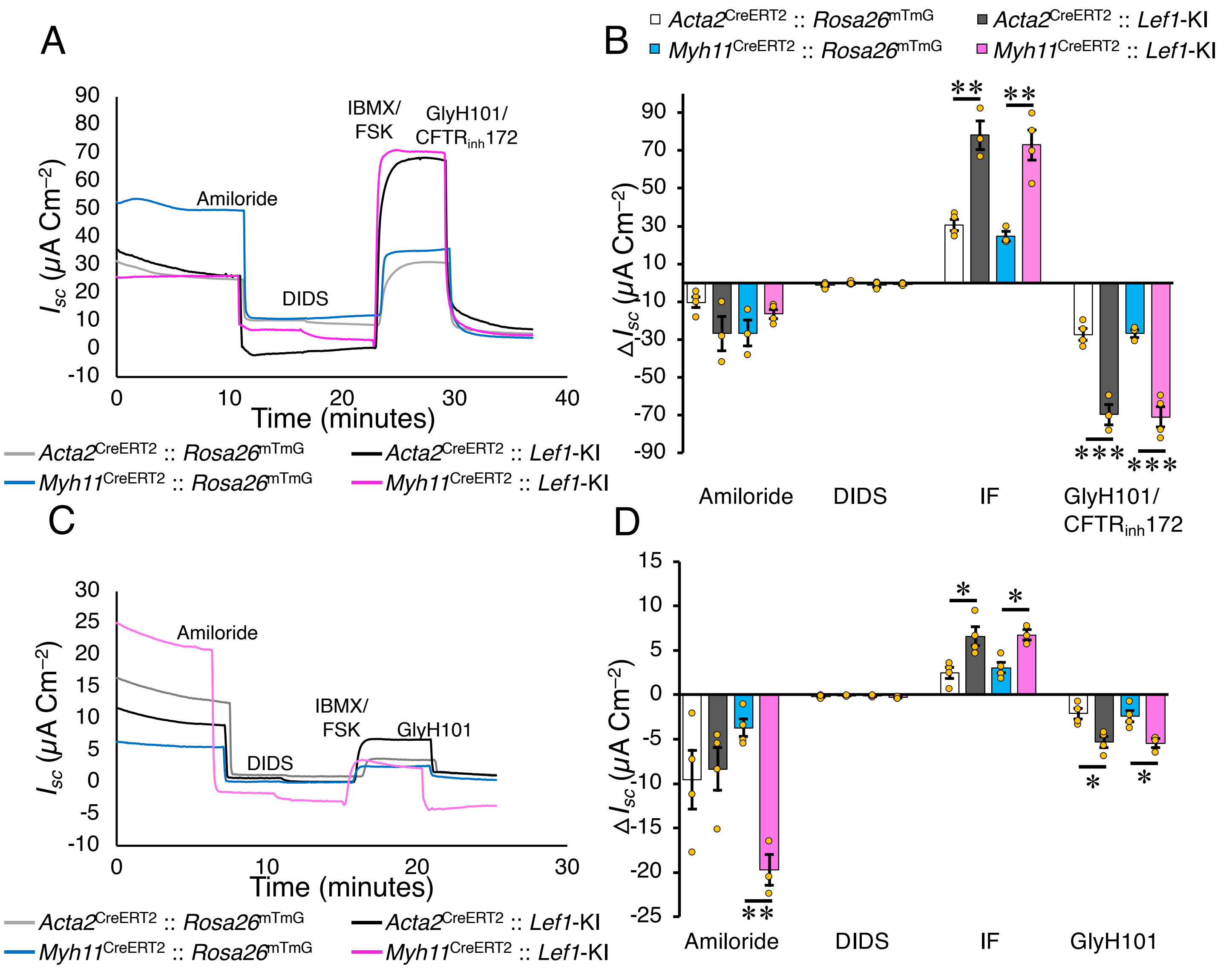


**Figure S2: Lef1 overexpression in GMECs enhances CFTR-mediated anion conductance.** Related to Figure 2. ALI membranes from the same experiments in Figure 3 underwent short circuit current measurement to assess chloride (**A-B**) and bicarbonate (**C-D**) conductance by measuring $I_{sc}$ on polarized GMEC grown at an air-liquid interface. Shown are traces of representative experiments (**A and C**) and the changes in $I_{sc}$ ($\Delta I_{sc}$) (**B and D**) in response to sequential addition of amiloride, 4,4′-diisothiocyanatostilbene2,2′-disulfonic acid (DIDS), forskolin + 3-isobutyl-1-methylxanthine (IBMX/FSK), and GlyH101 + $CFTR_{Inh}$ 172 (CFTR inhibitors), as indicated for the given genotypes. n= 4 donor groups. *$P$<0.05, **$P$<0.01, and ***$P$<0.001, two-way ANOVA followed by Bonferroni posttest. Error bars indicate SEM.

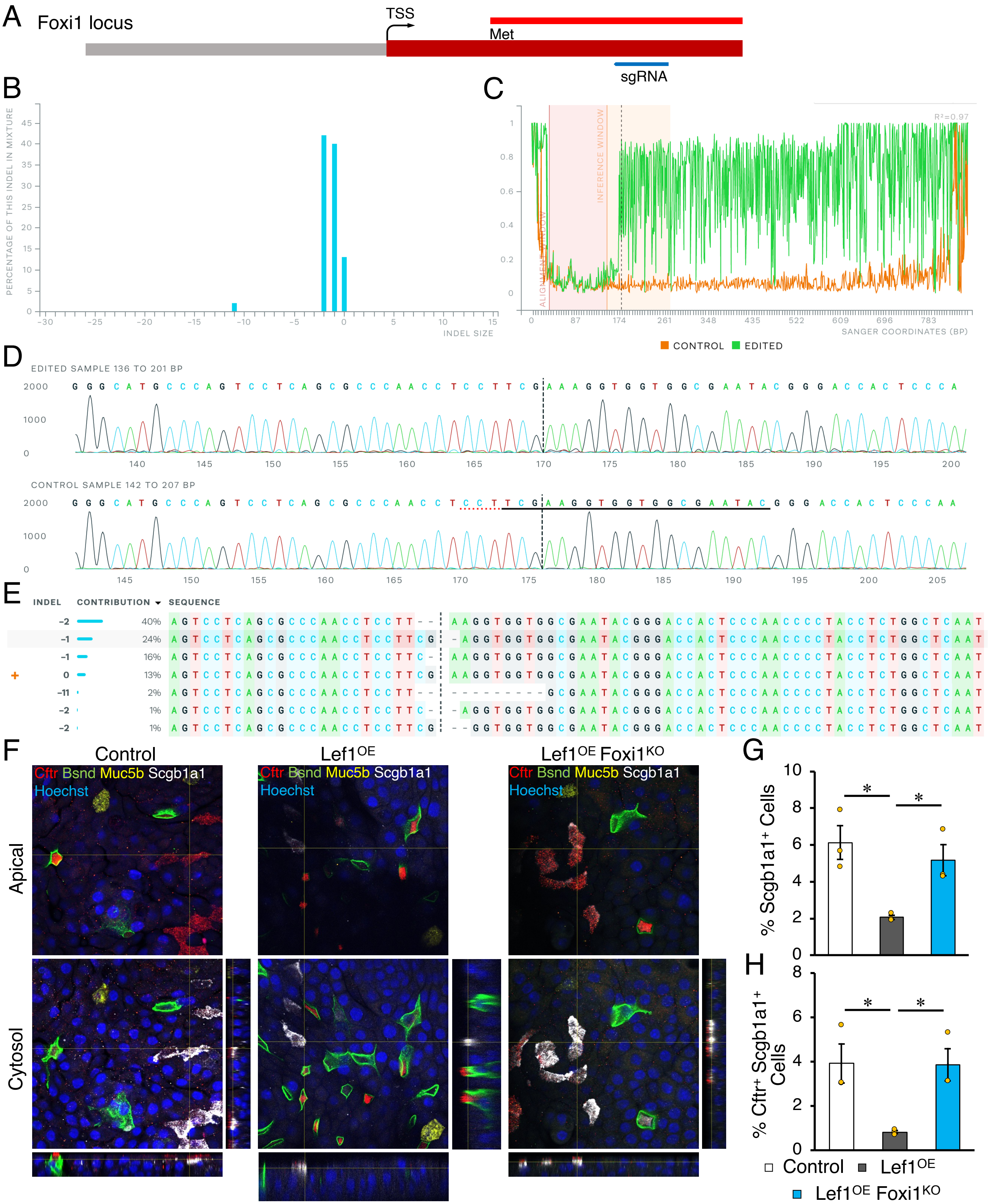


**Figure S3. Impact of *Foxi1* deletion on Lef1-mediated secretory specification.** Related to Figure 4. (**A**–**E**) Synthego ICE analysis of mutations found in target cells. (**A**) Schematic showing the location of sgRNA used for targeting at the *Foxi1* locus. TSS is transcription start site and Met is the first methionine. (**B**) Indel plot displaying the inferred distribution of indels at the *Foxi1* locus. This example shows 87% editing with out of frame indels. (**C**) A discordance plot detailing the level of alignment per base between the wild type control (in orange) and the edited sample (in green) within the inference window (the region around the cut site). It shows the average amount of signal that disagrees with the reference sequence derived from the control trace file. On the plot, the green line and orange line should be close together before the cut site, while these lines diverge near the cut site and continuing to remain far apart after the cut site (representing a high level of sequence discordance). (**D**) Sanger sequence of isolated clones showing edited and wild-type (control) sequences in the region around the guide sequence. This shows sequence base calls from both the control and the experimental sample. The horizontal black underlined region represents the guide sequence. The horizontal red dotted line is the PAM site. The vertical black dotted line represents the actual cut site. (**E**) Examples of indels present in the edited population and their relative proportions. Cut site is represented by black vertical dotted lines, and the wild-type sequence is marked by a "+" symbol on the far left. (**F**–**H**) Immunofluorescent images of polarized ALI cultures from the experiment in Figure 4. (**F**) *En face* and orthogonal images from whole mount ALI membranes derived from control, Lef1$^{OE}$, and Lef1$^{OE}$ Foxi1$^{KO}$ SMG progenitors immunolabelled for Cftr (red), Scgb1a1 (white), Bsnd (Green) and Muc5b (Yellow). (**G** and **H**) Quantification of Scgb1a1$^{+}$ (**G**) and Cftr$^{+}$ Scgb1a1$^{+}$ (**H**) cells. (n= 3 donor groups). **P*<0.05, one-way ANOVA followed by original FDR method of Benjamini and Hochberg posttest. Error bars indicate SEM.

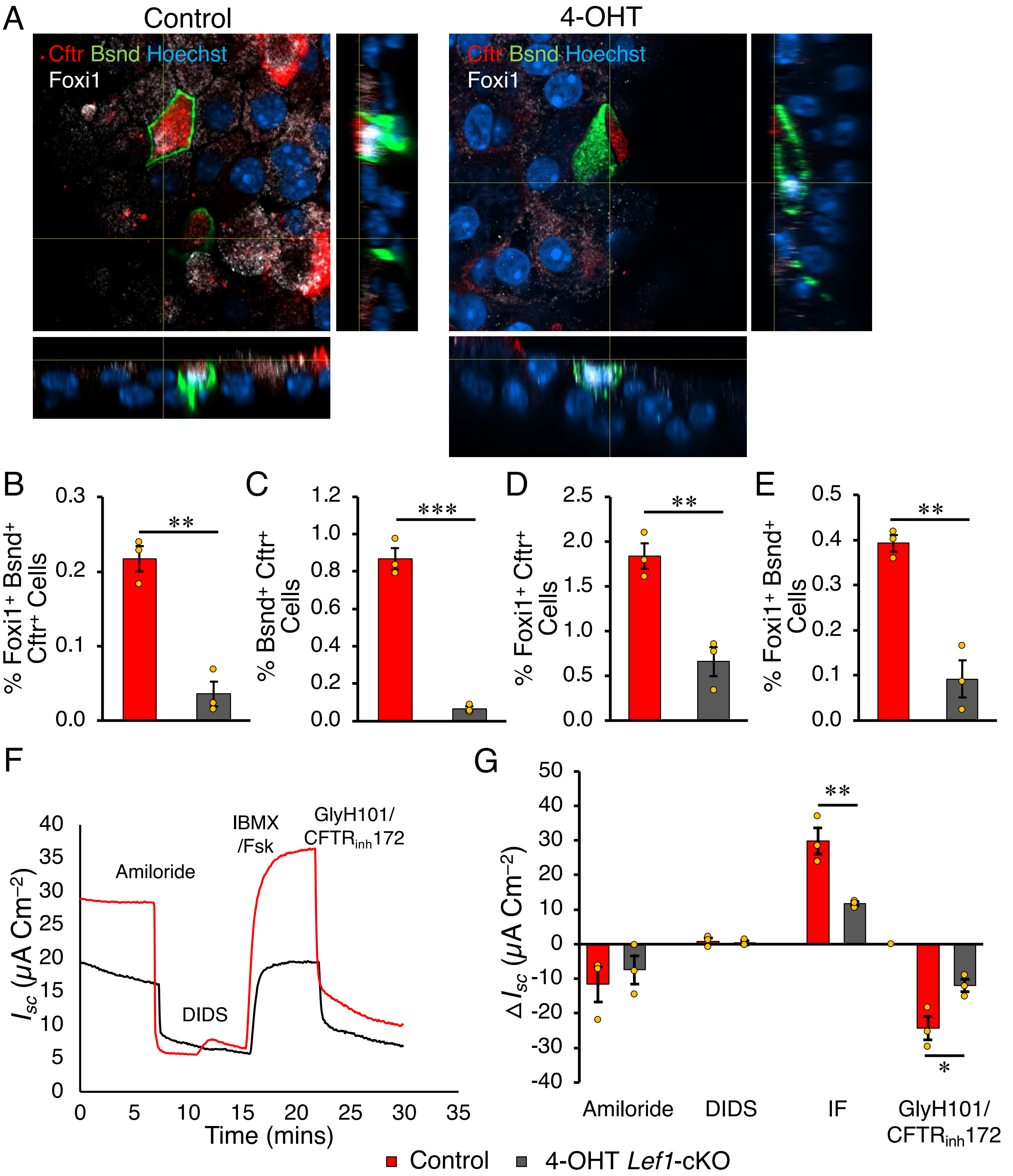


**Figure S4: Disruption of *Lef1* in SMG progenitors inhibits ionocyte specification and CFTR-mediated anion conductance.** Related to Figure 4. SMG progenitors were prepared from *Rosa26*$^{CreERT2}$::*Rosa26*$^{mTmG}$::*Lef1*-cKO triple transgenic mice and were plated on transwell inserts in SAGM + supplements for 3 days then maintained in PneumaCult ALI on the basolateral surface to establish an air liquid interface (ALI) culture. Half of the transwells were maintained in medium containing 2 µg/ml of 4-hydroxytamoxifen (4-OHT) for the entire length of the experiment. Six weeks later short circuit currents were assessed followed by fixation and wholemount immunofluorescence staining. (**A–E**) *En face* and orthogonal images of immunofluorescent whole mount membranes stained for Cftr (Red), Bsnd (Green) and Foxi1 (White) (**A**) and quantification of Foxi1$^+$ Bsnd$^+$ Cftr$^+$ (**B**), Bsnd$^+$ Cftr$^+$ (**C**), Foxi1$^+$ Cftr$^+$ (**D**) and Foxi1$^+$ Bsnd$^+$ (**E**) cells (n=3 donor groups). (**F–G**) Short-circuit current ($I_{sc}$) measurements under symmetrical chloride conditions. Shown are traces of representative experiments (**F**) and the change in $I_{sc}$ ($\Delta I_{sc}$) (**G**) in response to sequential addition of amiloride, 4,4′-diisothiocyanatostilbene 2,2′-disulfonic acid (DIDS), forskolin + 3-isobutyl-1-methylxanthine (IBMX/Fsk), and GlyH101. n= 3 donor groups. *$P$<0.05, **$P$<0.01, ***$P$<0.001 and ****$P$<0.0001, two-tailed Student's t test. Error bars indicate SEM.
